# Evaluation of *in vitro* activity of manogepix against multidrug-resistant and pan-resistant *Candida auris* from the New York Outbreak

**DOI:** 10.1101/2020.06.02.129916

**Authors:** YanChun Zhu, Shannon Kilburn, Mili Kapoor, Sudha Chaturvedi, Karen Joy Shaw, Vishnu Chaturvedi

## Abstract

An ongoing *Candida auris* outbreak in the New York metropolitan area is the largest recorded to date in North America. Laboratory surveillance revealed NY *C. auris* isolates are resistant to fluconazole, with variable resistance to other currently used broad-spectrum antifungal drugs, and that several isolates are pan-resistant. Thus, there is an urgent need for new drugs with a novel mechanism of action to combat the resistance challenge. Manogepix (MGX) is a first-in-class agent that targets the fungal Gwt1 enzyme. The prodrug, fosmanogepix, is currently in Phase 2 clinical development for the treatment of fungal infections. We evaluated the susceptibility of 200 New York *C. auris* isolates to MGX and 10 comparator drugs using CLSI methodology. MGX demonstrated lower MICs than comparators (MIC_50_ and MIC_90_ 0.03 mg/L; range 0.004-0.06 mg/L). The MGX epidemiological cutoff value (ECV, 99% cutoff) for the tested *C. auris* isolates was 0.06 mg/L. MGX was 8-32-fold more active than the echinocandins, 16-64-fold more active than the azoles, and 64-fold more active than amphotericin B. No differences were found in the MGX or comparators’ MIC_50_, MIC_90_, or GEOMEAN values when subsets of clinical, surveillance, and environmental isolates were evaluated. The range of MGX MIC values for six *C. auris* pan-resistant isolates was 0.008-0.015 mg/L, and the median and mode MIC values were 0.015 mg/L, demonstrating that MGX retains activity against these isolates. These data support further clinical evaluation of fosmanogepix for the treatment of *C. auris* infections, including highly resistant isolates.

## INTRODUCTION

The yeast pathogen *Candida auris* has been shown to be responsible for severe illnesses among hospitalized patients in the New York metropolitan area (1). Of 1, 092 US *C. auris* clinical cases tracked by the CDC, 661 (60.53%) were reported from New York and New Jersey (https://www.cdc.gov/fungal/candida-auris/tracking-c-auris.html). *C. auris* cases in New York are concentrated among hospitalized patients and nursing home residents, with 504 clinical cases and 747 screening cases confirmed as of May 15, 2020 (https://www.health.ny.gov/diseases/communicable/c_auris/). An earlier analysis of New York *C. auris* cases indicated 23 (45%) of the 51 clinical case-patients died within 90 days (1, 2). Initial reports suggested that *C. auris* emerged independently in four regions (South Asia, East Asia, Africa and South America) defined by clades I, II, II and IV, respectively (3) with Iranian clade V arising more recently (4). Analysis of a diverse collection of *C. auris* isolates showed that clade I had the highest percentage of fluconazole resistant, amphotericin B resistant as well as MDR and XDR isolates (5). *C. auris* isolates in New York are resistant to fluconazole and many strains also demonstrated elevated MICs to voriconazole (VRC), amphotericin B (AMB), flucytosine (5FC), and echinocandins, consistent with the finding that they were largely members of clade I (6). More recently, pan-resistant *C. auris* isolates (resistant to two or more azoles, AMB, and echinocandins) were recorded in New York (7). Clinicians and public health professionals face serious challenges in dealing with the large, sustained outbreak of *C. auris* in New York that is exacerbated by the limited availability of effective treatment options (8, 9). Thus, there is an urgent need to evaluate the efficacy of new or re-purposed antifungal drugs against drug-resistant *C. auris*.

Manogepix (MGX, APX001A) is a first-in-class small molecule antifungal agent that targets the fungal inositol acylase (Gwt1), which catalyzes an early step in the glycosylphosphatidyl inositol (GPI)-anchor biosynthesis pathway (10–12). Inhibition of Gwt1 prevents the appropriate localization of cell wall mannoproteins, which compromises cell wall integrity, biofilm formation, germ tube formation and results in severe fungal growth defects (13, 14). The phosphatidylinositol glycan anchor biosynthesis class W (PIGW) protein is the closest mammalian ortholog of the fungal Gwt1 protein and is not inhibited by MGX (13). The N-phosphonooxymethyl prodrug, fosmanogepix, is currently in clinical trials for the treatment of invasive fungal infections caused by *Candida* including *C. auris*, Aspergillus and rare molds (15, 16).

Several studies have previously evaluated the activity of MGX against world-wide or Indian collections of *C. auris* strains using both CLSI and EUCAST methodologies (17–20). MGX also demonstrates excellent activity against other *Candida* spp, with the exception of *C. krusei*, as well as activity against *Aspergillus, Scedosporium, Fusarium, Lomentospora* and members of the Mucorales order (21–23). In addition, MGX retains activity against AMB- and azole-resistant strains of *Aspergillus* spp. and azole- and echinocandin-resistant *Candida* spp. (22–24). Efficacy studies in a wide range of yeast and mold species demonstrated significant improvements in both survival as well as colony forming unit (CFU) endpoints in lung, kidney, and brain, consistent with wide tissue distribution (18, 24–27).

The present study describes the *in vitro* activity of MGX and 10 comparators against a collection of 200 clinical, environmental, and surveillance isolates of *C. auris* collected in New York between 2017-2020, with the majority (94%) of isolates collected between 2018 and 2019 (6).

## RESULTS

The MGX antifungal susceptibility profile of NY *C. auris* isolates in comparison with 10 other antifungals, is summarized in Tables 1-2. All 200 NY *C. auris* isolates were classified as resistant to fluconazole (FLC) with MIC values ≥32 mg/L. A comparison of MIC_90_ values show that MGX was 8-32-fold more active than the echinocandins anidulafungin (AFG), caspofungin (CAS), and micafungin (MFG); 16-64 fold more active than the azoles isavuconazole (ISA), itraconazole (ITC), posaconazole (POS) and VRC; 64-fold more active than AMB, and >1000-fold more active than 5FC (Table 1).

**Table 1.** Summary of Susceptibility (mg/L) of 200 NY *C. auris* isolates

| Antifungal | GEOMEAN | MODE | MIC<sub>50</sub> | MIC<sub>90</sub> | RANGE |
| --- | --- | --- | --- | --- | --- |
| MGX | 0.02 | 0.03 | 0.03 | 0.03 | 0.004 – 0.06 |
| AFG | 0.33 | 0.50 | 0.25 | 1.0 | 0.03 – 8 |
| CAS | 0.14 | 0.25 | 0.12 | 0.25 | 0.016 – >16 |
| MFG | 0.16 | 0.12 | 0.12 | 0.25 | 0.06 – 4 |
| FLC | 246.42 | 256 | 256 | 256 | 32 – >256 |
| ISA | 0.71 | 1 | 1 | 1.0 | 0.03 – 2 |
| ITC | 0.58 | 1 | 0.5 | 1.0 | 0.125 – 1 |
| POS | 0.20 | 0.25 | 0.25 | 0.5 | 0.03 – 1 |
| VRC | 1.67 | 2 | 2 | 2 | 0.06 – 4 |
| AMB | 1.35 | 2 | 1 | 2 | 0.125 – 32 |
| 5FC | 0.14 | 0.06 | 0.064 | 32 | 0.023 – 32 |

**Table 2.**
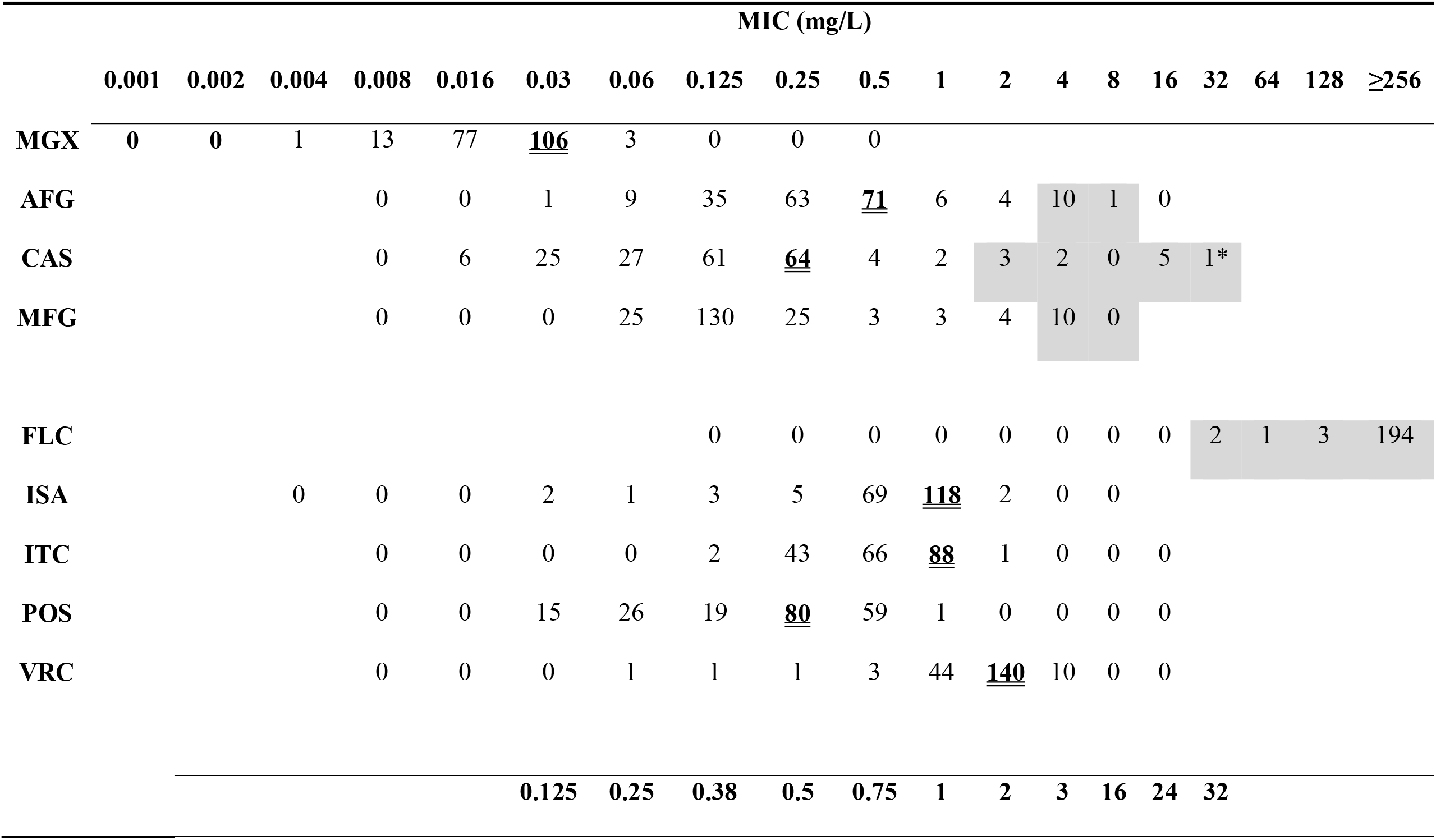
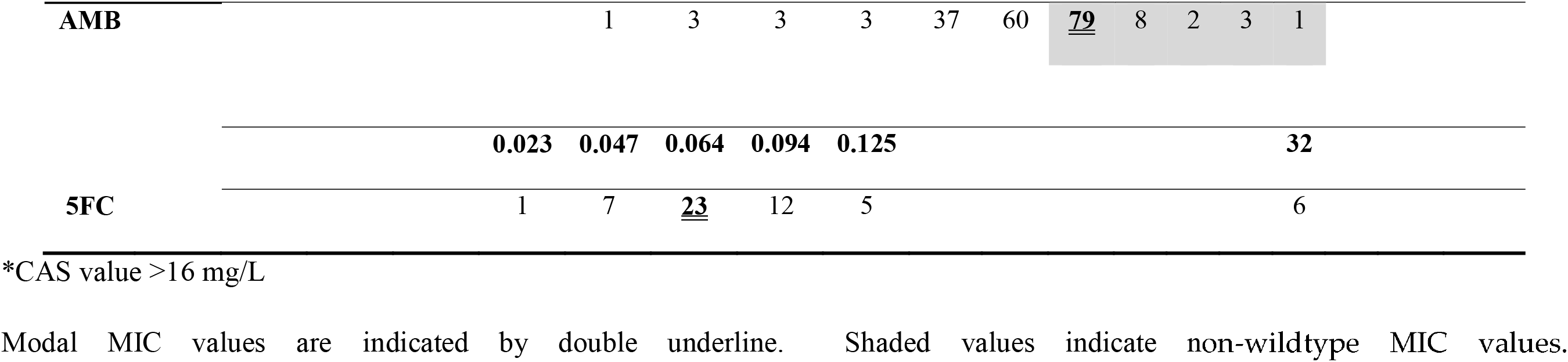
*C. auris* MIC distributions for MGX and Comparators

|  | MIC (mg/L) |  |  |  |  |  |  |  |  |  |  |  |  |  |  |  |  |  |  |
| --- | --- | --- | --- | --- | --- | --- | --- | --- | --- | --- | --- | --- | --- | --- | --- | --- | --- | --- | --- |
|  | 0.001 | 0.002 | 0.004 | 0.008 | 0.016 | 0.03 | 0.06 | 0.125 | 0.25 | 0.5 | 1 | 2 | 4 | 8 | 16 | 32 | 64 | 128 | ≥256 |
| MGX | 0 | 0 | 1 | 13 | 77 | <u>106</u> | 3 | 0 | 0 | 0 |  |  |  |  |  |  |  |  |  |
| AFG |  |  |  | 0 | 0 | 1 | 9 | 35 | 63 | <u>71</u> | 6 | 4 | 10 | 1 | 0 |  |  |  |  |
| CAS |  |  |  | 0 | 6 | 25 | 27 | 61 | <u>64</u> | 4 | 2 | 3 | 2 | 0 | 5 | 1* |  |  |  |
| MFG |  |  |  | 0 | 0 | 0 | 25 | 130 | 25 | 3 | 3 | 4 | 10 | 0 |  |  |  |  |  |
| FLC |  |  |  |  |  |  |  | 0 | 0 | 0 | 0 | 0 | 0 | 0 | 0 | 2 | 1 | 3 | 194 |
| ISA |  |  | 0 | 0 | 0 | 2 | 1 | 3 | 5 | 69 | <u>118</u> | 2 | 0 | 0 |  |  |  |  |  |
| ITC |  |  |  | 0 | 0 | 0 | 0 | 2 | 43 | 66 | <u>88</u> | 1 | 0 | 0 | 0 |  |  |  |  |
| POS |  |  |  | 0 | 0 | 15 | 26 | 19 | <u>80</u> | 59 | 1 | 0 | 0 | 0 | 0 |  |  |  |  |
| VRC |  |  |  | 0 | 0 | 0 | 1 | 1 | 1 | 3 | 44 | <u>140</u> | 10 | 0 | 0 |  |  |  |  |
|  |  |  |  |  |  | <u>0.125</u> | <u>0.25</u> | <u>0.38</u> | <u>0.5</u> | <u>0.75</u> | <u>1</u> | <u>2</u> | <u>3</u> | <u>16</u> | <u>24</u> | <u>32</u> |  |  |  |

|  |  |  |  |  |  |  |  |  |  |  |  |
| --- | --- | --- | --- | --- | --- | --- | --- | --- | --- | --- | --- |
| **AMB** | 1 | 3 | 3 | 3 | 37 | 60 | <u>**79**</u> | 8 | 2 | 3 | 1 |
|  | **0.023** | **0.047** | **0.064** | **0.094** | **0.125** |  |  |  |  |  | **32** |
| **5FC** | 1 | 7 | <u>**23**</u> | 12 | 5 |  |  |  |  |  | 6 |

\*CAS value >16 mg/L
Modal MIC values are indicated by double underline. Shaded values indicate non-wildtype MIC values.

MGX demonstrated a narrow distribution range (within 5 two-fold serial dilutions) and the MIC_50_, MIC_90_ and modal MIC values were 0.03 mg/L (Table 2). When the subsets of clinical (e.g, blood, abscess, wound), surveillance (e.g., nares, groin, axilla) and environmental isolates were evaluated, no differences (within 1-2 fold) were found in the MGX or comparators’ MIC_50_, MIC_90_, or GEOMEAN values (Supplementary Table 1, Supplementary Table 2). The epidemiologic cutoff value (ECV) 99 % was determined using the dataset of 200 isolates as described for ECOFFinder XL 2010 v2.1 (Figure 1)(28). The MGX ECV was 0.06 mg/L, suggesting that all isolates were within the population of wildtype (WT) strains.

**Figure 1.**
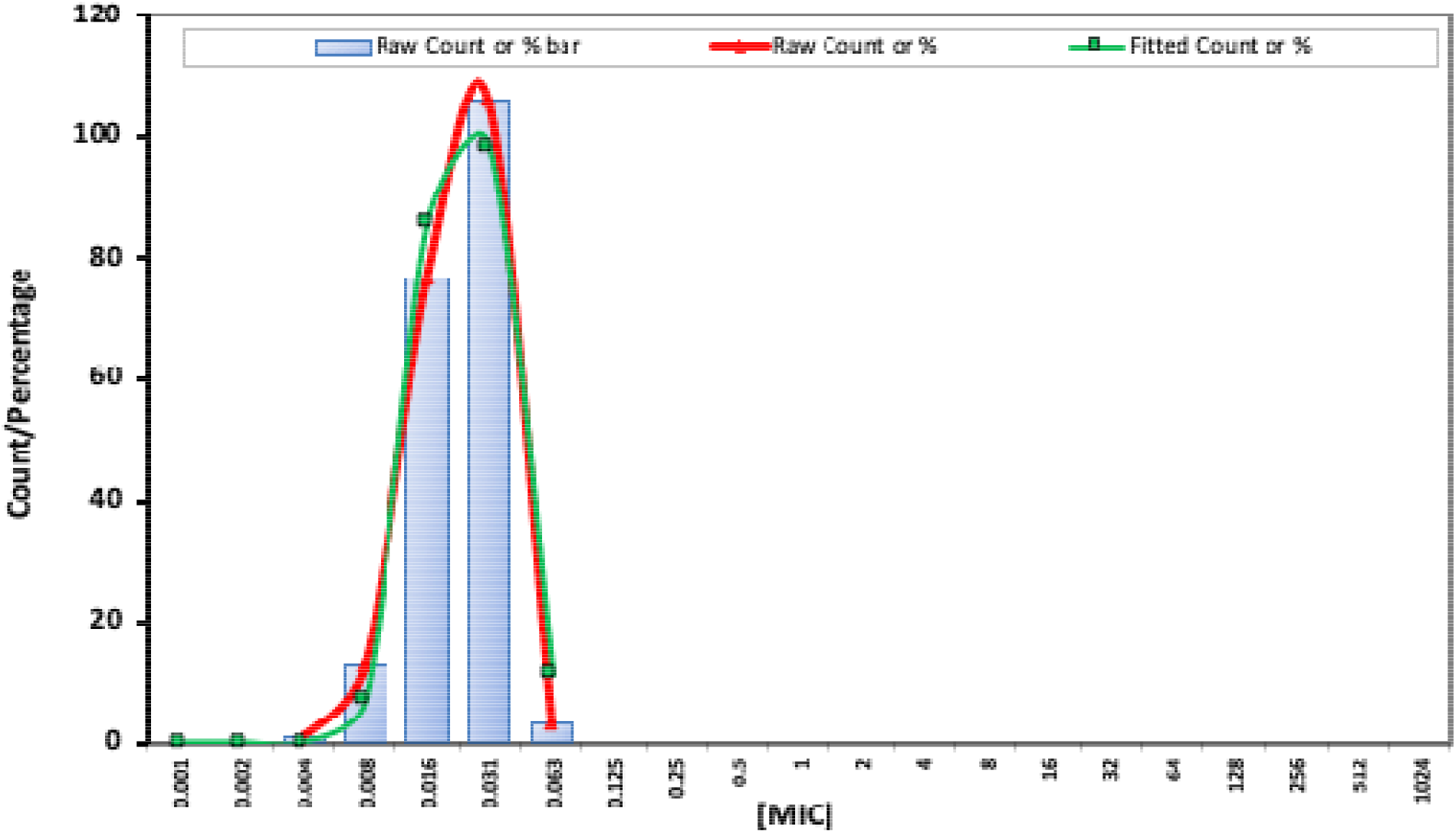
Epidemiological cutoff values (ECVs) of MGX for NY *C. auris* isolates determined using ECOFFinder.

Limited 5FC susceptibility testing of 54 isolates also demonstrated a narrow MIC range (0.023 to 0.125 mg/L), with the exception of 6 non-susceptible isolates demonstrating MIC values of 32 mg/L (Table 2). The range of values for the echinocandins spanned 9, 11, and 8, two-fold serial dilutions for AFG, CAS and MFG, respectively (Table 2).

Non-wildtype MIC values were evaluated, based upon CLSI susceptibility breakpoints, CDC recommendations for susceptibility breakpoints, and published *C. auris* ECVs [AMB ≥2 mg/L, AFG ≥4 mg/L, CAS 2 mg/L, and MFG ≥4 mg/L]((19, 29) https://www.cdc.gov/fungal/candida-auris/c-auris-antifungal.html). A total of 13 *C. auris* isolates demonstrated MIC values greater than or equal to these values for one or more of the echinocandins, with 7 isolates demonstrating evidence of cross-resistance to all 3 echinocandins (Supplementary Table 3). GEOMEAN values for these 13 isolates were 3.79 mg/L (AFG), 4.76 mg/L (CAS), and 3.06 mg/L (MFG). Of note, the GEOMEAN for MGX vs this collection of isolates was 0.013 mg/L, which is less than the GEOMEAN for the entire collection of 200 isolates (0.02 mg/L), consistent with the lack of cross-resistance to the echinocandins.

The MIC values for AMB, as determined by ETEST® (bioMérieux), ranged from 0.125 to 32 mg/L and the GEOMEAN, MODE, MIC_50_ and MIC_90_ values were 1.35, 2, 1, and 2 mg/L, respectively (Supplementary Table 4). The wide range of values obtained (9 two-fold serial dilutions) suggested non-WT isolates within the collection. Based upon the previously determined *C. auris* AMB ECV of AMB ≥2 mg/L, >50% of the population in this study was non-WT. Using a more stringent cutoff of 3 mg/L resulted in the identification of 14 isolates (7% of the total study population) with potentially non-WT AMB MIC values, with 6 isolates demonstrating AMB MIC values ≥16 mg/L (Supplementary Table 4). None of the isolates in Supplementary Table 3 (echinocandin non-susceptible) were within this group of 14 isolates and the GEOMEAN values for AFG, CAS and MFG were lower for the collection of 14 isolates than the larger collection of 200 isolates. MGX GEOMEAN MIC values for the total collection of 200 isolates vs the 14 isolates were 0.22 vs 0.26 mg/L. These data are consistent with a lack of cross-resistance between AMB and the echinocandins as well as AMB and MGX.

The azoles as a group showed a narrower range of MIC values (5-7 two-fold serial dilutions) than either the echinocandins or AMB (Table 2). All *C. auris* isolates were FLC-resistant using the tentative breakpoint of ≥32 mg/L. Resistance categories have not been established for *C. auris* for ISA, ITC, POS and VRC; however for *C. albicans*, isolates are classified as FLC resistant (≥8 mg/L) or VRC resistant (≥0.5 mg/L EUCAST) whereas for *C. parapsilosis*, VRC resistant ERG11 mutants were identified with MIC values of ≥1 mg/L (30–33). Twelve strains were identified with increased azole MIC values of which 10 strains had VRC MIC values of 4 mg/L, and an additional two strains had both ISA and VRC MIC values of 2 mg/L (Supplementary Table 5). Tentative epidemiological cutoff values for *C. auris* using CLSI reference methods have been evaluated and ECOFFs (using ECOFF finder) for 95%/99% endpoints were: ITC, 0.5/1; VRC, 8/32; ISA, 1/2; POS, 0.125/0.25 (19). Using the 99% endpoint values, the data in Table 2 suggest that the numbers of isolates that are non-WT are: ITC, 1; VRC, 0; ISA, 0; POS, 60. Using the 95% endpoint values, the data in Table 2 suggest that the numbers of isolates that are non-WT are: ITC, 89; VRC, 0; ISA, 2; POS 140 isolates. Using the rule of thumb that the ECOFF is two-fold dilutions higher than the mode would suggest that none of the isolates are resistant to any of the azoles, other than FLC.

Five NY *C. auris* isolates were previously identified as pan-resistant, based upon the definition of elevated MIC values to FLC, AMB and the echinocandins (6, 7, 34). The MGX susceptibility of these isolates, plus a sixth isolate obtained in 2020 are shown in Table 3. All six pan-resistant *C. auris* isolates were susceptible to MGX (0.008-0.016 mg/L) and had MIC values below the modal value of the larger collection.

**Table 3.** MGX MIC values for pan-resistant *C. auris*

| *C. auris* |  | MIC (mg/L) |  |  |  |  |  |  |  |  |  |
| --- | --- | --- | --- | --- | --- | --- | --- | --- | --- | --- | --- |
| strain | FLC | VRC | ITC | ISA | POS | AFG | CAS | MFG | AMB | 5FC | MGX |
| 17-24 | 256 | 2 | 0.5 | 0.5 | 0.25 | 4 | >16 | 4 | 2 | 0.064 | 0.015 |
| 18-2 | 256 | 2 | 1 | 1 | 0.25 | 8 | 2 | 4 | 2 | 0.064 | 0.015 |
| 19-42 | 256 | 2 | 1 | 1 | 0.25 | 4 | 16 | 4 | 2 |  | 0.015 |
| 19-43 | 256 | 2 | 1 | 1 | 0.25 | 4 | 16 | 4 | 2 |  | 0.015 |
| 19-4 | 256 | 2 | 0.5 | 0.5 | 0.25 | 4 | 2 | 4 | 2 |  | 0.015 |
| 20-1 | 256 | 2 | 0.5 | 1 | 0.25 | 4 | 4 | 4 | 2 |  | 0.007 |

Clade information was available for 62 of the 200 isolates. All were identified as South Asia clade 1, consistent with previous observations that NY *C. auris* isolates largely belong to South Asia clade I (6).

## DISCUSSION

*C. auris* is an urgent threat in the United States (CDC’s Antibiotic Resistance Threats in the United States, 2019 https://www.cdc.gov/drugresistance/biggest-threats.html). *C. auris* isolates involved in the largest US outbreak in the New York metropolitan area are resistant to FLC, variably resistant to other antifungal drugs, and an emergent danger due to observed pan-resistance (6, 7, 34). *C. auris* isolates previously recovered in New York were 100% resistant to FLC, 81% resistant to VRC (MIC >2.0 mg/ L), 61% to AMB (MIC >2.0 mg/ L), with emerging resistance to one or more echinocandins (6). The observed pattern is higher than reported in other published studies including South Asian clade I isolates (3, 29, 35), and also included organisms with AMB MIC values as high as 32 mg/L. Although the underlying mechanisms remain partially unknown, active laboratory surveillance of *C. auris* antifungal resistance patterns as well as mechanism identification remain high priorities. There is an urgent need for the development of new antifungal drugs with different mechanisms of action against *C. auris* to manage this serious resistance challenge. Additionally, diagnostic laboratories need to expand antifungal test panels to include newly licensed drugs in order to provide evidence-based input for the clinical and public health management of drug-resistant *C. auris*.

MGX was the most active antifungal agent tested against the 200 *C. auris* isolates in the present study and this activity was independent of the clinical, environmental or surveillance source of *C. auris* (Supplementary Table 2). Similarly, no differences in the spectrum of activity of the isolates in the 3 subsets was observed for the comparator drugs, suggesting that the isolates may be derived from common sources. The laboratory-observed pattern of resistance to comparator antifungals did not impact MGX *in vitro* activity in the present study. The MGX MIC range 0.004 – 0.06 mg/L was within two dilutions of the ranges reported earlier (17, 18). Additionally, MGX ECV (0.06 mg/L) in the present study was within two-dilutions of 0.016 mg/L Wild-Type upper limit (WT-UL) reported for a collection of 122 Indian *C. auris* isolates (South Asia clade I) evaluated by EUCAST methodology (19).

MGX, the active moiety of fosmanogepix (APX001), shows potent activity against various *Candida* species except *C. krusei*, and a wide variety of pathogenic molds (21, 22). The pharmacokinetic (PK)/pharmacodynamic (PD) evaluation of fosmanogepix in the neutropenic mouse model of disseminated candidiasis demonstrated concentration-dependent in vivo efficacy against *C. auris* and other *Candida* species (24). In a *C. auris* disseminated model of infection, mice treated with fosmanogepix demonstrated higher rate of survival vs AFG (18). Similarly, in a delayed treatment model utilizing a fluconazole-resistant strain, fosmanogepix-treated mice demonstrated significant improvements in survival at day 21 as well as reductions in kidney burden (20). The fosmanogepix efficacy observed in animal models awaits further confirmation from ongoing trials for invasive candidiasis/candidemia (36).

The promising profile of MGX includes a unique mode of action and low potential for the development of spontaneous resistance for several *Candida* species (37). Further examination of the spontaneous mutants of *C. albicans* and *C. parapsilosis* by whole genome sequencing and mutant analysis revealed two efflux-mediated mechanisms, which lead to a 4-8-fold increase in MGX MIC values and a concomitant 2-4-fold increase in FLC MIC values (38). Previously, a link between elevated MGX values in a subset of FLC resistant strains was suggested possibly due to efflux-mediated mechanisms in *C. auris* as well as other pathogenic yeasts (19, 39). For *C. auris*, Arendrup et al. (19) reported a wider range of MGX (EUCAST) MIC values (range 0.001-0.125 mg/L; mode 0.016 mg/L) than observed here. Further analysis showed that isolates with high (> 64 mg/L) FLC MICs had a higher range of MGX MICs (0.004-0.06 mg/L) whereas those with lower FLC MIC values (≤ 64 mg/L) demonstrated a lower range of MGX MICs (0.001-0.008 mg/L) with MGX MIC values of 0.004 or 008 mg/L found in both groups. Although elevated MGX MICs were associated with higher FLC MICs not all of the higher FLC MICs demonstrated elevated MGX MICs suggesting that some, but not all, FLC resistance mechanisms affect MGX susceptibility (19). In the current study, where all of the isolates had FLC MIC values of ≥256 mg/L, a narrower range of MGX MICs was observed (0.004-0.06 mg/L) consistent with the range of values observed in the Arendrup et al study (19). However, although the range of MGX MIC values may be increased 2 to 4-fold in some highly FLC resistant strains, the MGX MIC values remain low and the clinical impact on the efficacy of fosmanogepix is as yet unknown.

Our observations revealed (1) low MGX MICs against the largest collection of *C. auris* isolates tested to date, (2) MGX MICs were unchanged against *C. auris* isolates with variable pattern of resistance to amphotericin B, azoles, and echinocandins, (3) MGX retained potent *in vitro* activity against six pan-resistant *C. auris* isolates, (4) the CLSI broth microdilution method was reproducible for MGX testing of *C. auris*. The potent activity of this novel antifungal agent against *C. auris* supports the continued clinical evaluation for fosmanogepix in the treatment of these often-resistant infections. These findings would also allow us to expand the antifungal susceptibility testing panel to include MGX evaluation of *C. auris* from NY metropolitan area, which might help guide evidence-based management of *C. auris* clinical and surveillance cases.

## MATERIALS AND METHODS

### Fungal isolates

The details of methods used for processing of the samples for the isolation and characterization *C. auris* were described recently (6). Two hundred *C. auris* isolates were selected at random and consisted of 85 clinical, 97 surveillance, and 18 environmental isolates. The details of the various sample types yielding *C. auris* isolates, are summarized in Supplementary Table 1. The number of isolates evaluated from each year was as follows: 2017 (2), 2018 (58), 2019 (130), and 2020 (10).

### Antifungal susceptibility testing

The antifungals tested were FLC, VRC, ITC, ISA, POS, AFG, CAS, MFG, AMB and 5FC. Broth micro-dilution antifungal susceptibility testing was performed in accordance with Reference Method M60 of the Clinical and Laboratory Standards Institute using TREK frozen broth microdilution panels (catalog number CML2FCAN; Thermo Fisher Scientific, Marietta, OH, USA) for FLC, VRC, ITC, ISA, POS, AFG, CAS, and MFG (40). AMB and FLC MICs were determined by ETEST^®^ at 24 h (AB Biodisk; bioMérieux, Solna, Sweden). *C. albicans* ATCC 90028 and *C. parapsilosis* ATCC 22019, obtained earlier from the American Type Culture Collection, were used as quality control isolates. The susceptibility breakpoints of the Centers for Disease Control and Prevention were used to assess antifungal resistance patterns in *C. auris* (FLC ≥32, AMB ≥2, AFG ≥4, CAS ≥2 and MFG≥4), and the ETEST^®^ AMB MIC value of 1.5 mg/L was rounded up to 2.0 mg/L as described (https://www.cdc.gov/fungal/candida-auris/c-auris-antifungal.html). Susceptibility/resistance to VRC and other triazoles (ITA, ISA, POS) were assessed using published reports for other *Candida* species (30–33). We also considered a prior publication on the epidemiological cut-off values of antifungal compounds for 122 *C. auris* isolates from India (19). As the term is not used conventionally with antifungal drugs, we defined pan-resistant *C. auris* as isolates with *in vitro* resistance to two or more azoles, all echinocandins, and AMB (34).

### Manogepix (MGX) susceptibility testing

MGX stock solutions were prepared at 10 mg/mL in 100% dimethyl sulfoxide (DMSO) and aliquots stored at −20 °C. Broth microdilution susceptibility testing was performed according to CLSI M27-A3 guidelines. MGX was first diluted in DMSO to obtain intermediate dilutions. These were further diluted in microtiter plates to obtain a final concentration of 0.001 to 0.5 mg/L. 1 μl of DMSO was added to “No drug” control wells. The solutions were mixed on a plate shaker for 10 mins and plates incubated at 35°C for 24 h. For MGX, the minimum concentration that led to 50% reduction in fungal growth as compared to the control was determined as the minimum inhibitory concentration (MIC), as previously described for the echinocandins (41). *C. albicans* ATCC 90028 and *C. parapsilosis* ATCC 22019, obtained earlier from the American Type Culture Collection, were used as quality control isolates. MGX was tested seven times against two QC strains, and MIC values for *C. albicans* ATCC90028 and *C. parapsilosis* ATCC22019, were within recommended CLSI ranges (Supplementary Table 6).

### Epidemiology cutoff values (ECV)

The epidemiological cutoff values (ECVs, ECOFFs) for MGX were estimated using the Microsoft Excel spreadsheet calculator ECOFFinder (28).

## ACKNOWLEDGEMENTS

This study was supported by a grant from Amplyx Pharmaceuticals. M.K. is an employee of Amplyx; K.J.S., was a previous employee of Amplyx and is now an independent consultant for Amplyx. All other authors have no conflicts.

## SUPPLEMENTARY INFORMATION

**Supplementary Table 1.**
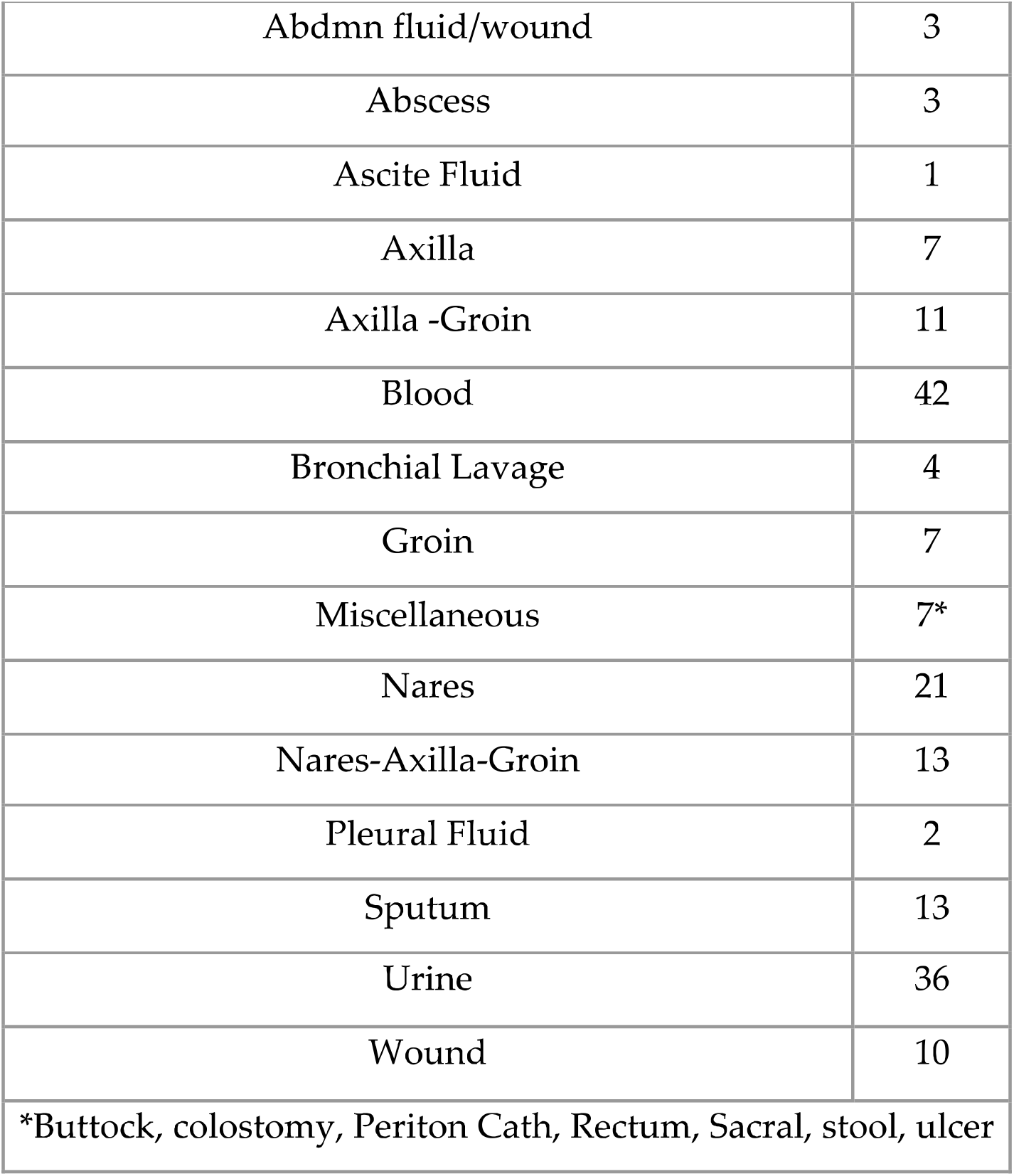
Sources and numbers of *C. auris* isolates included in the present study

**Supplementary Table 2.**
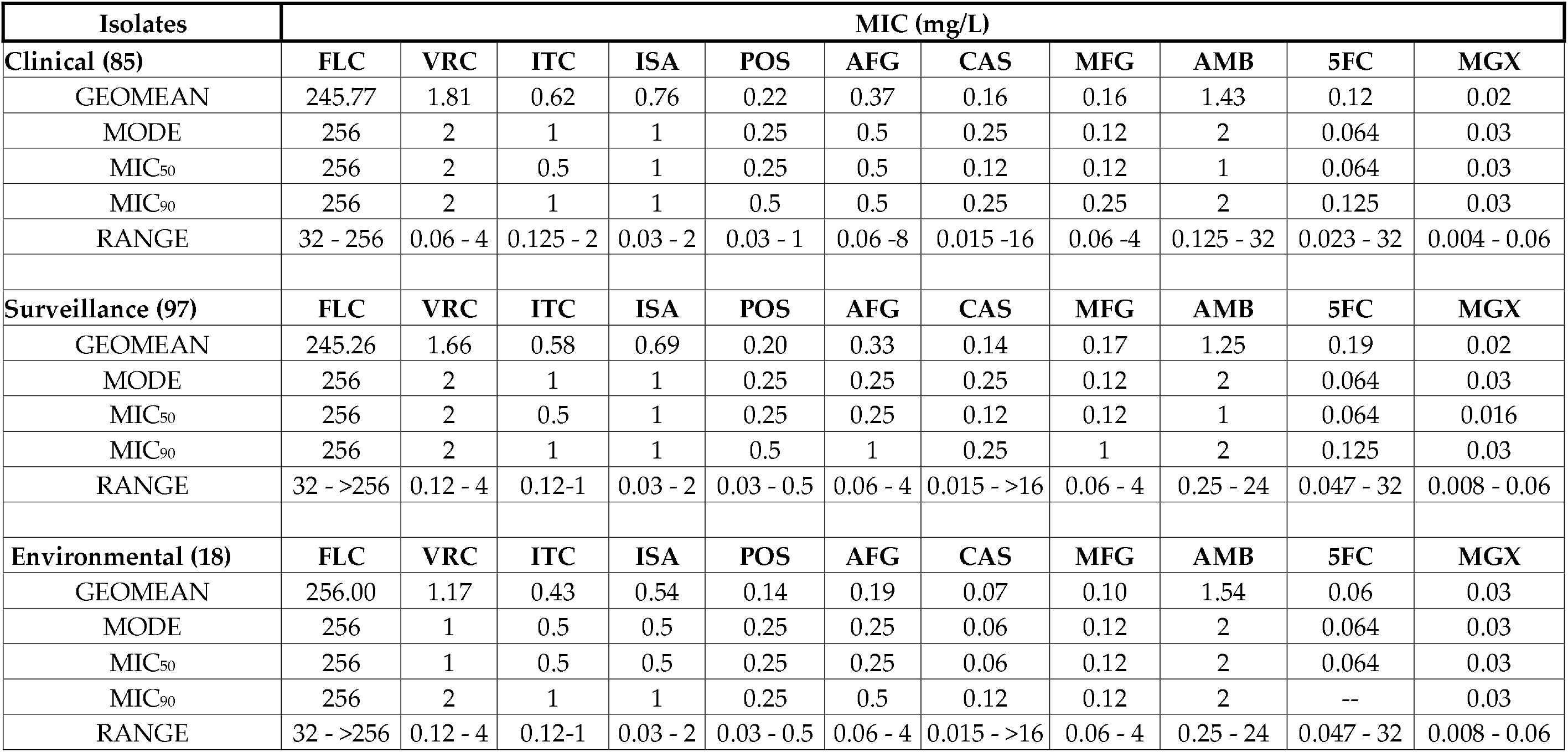
Susceptibility of antifungal agents in clinical, surveillance and environmental *C. auris* isolates

**Supplementary Table 3.** MGX MICs (mg/L) for *C. auris* isolates with high echinocandin MICs

| *C. auris* | AFG | CAS | MFG | MGX |
| --- | --- | --- | --- | --- |
| 17-24 | 4 | >16 | 4 | 0.015 |
| 18-2 | 8 | 2 | 4 | 0.015 |
| 19-42 | 4 | 16 | 4 | 0.015 |
| 19-43 | 4 | 16 | 4 | 0.015 |
| 19-4 | 4 | 2 | 4 | 0.015 |
| 20-1 | 4 | 4 | 4 | 0.008 |
| 17-1 | 4 | 4 | 4 | 0.015 |
| 18-17 | 2 | 1 | 4 | 0.015 |
| 19-26 | 2 | 1 | 4 | 0.015 |
| 19-44 | 4 | 16 | 1 | 0.008 |
| 19-45 | 4 | 16 | 1 | 0.008 |
| 19-22 | 4 | 16 | 2 | 0.008 |
| 19-128 | 4 | 2 | 4 | 0.015 |
| GEOMEAN | 3.792 | 4.757 | 3.064 | 0.0126 |

**Supplementary Table 4.** MGX MICs (mg/L) for 14 *C. auris* isolates with high AMB MICs (> 3.0 mg/L)

| *C. auris* | AMB | MGX |
| --- | --- | --- |
| 18-51 | 3 | 0.03 |
| 18-19 | 3 | 0.015 |
| 18-9 | 3 | 0.03 |
| 18-3 | 3 | 0.03 |
| 19-10 | 3 | 0.03 |
| 19-29 | 3 | 0.03 |
| 19-32 | 3 | 0.03 |
| 19-129 | 3 | 0.015 |
| 19-130 | 16 | 0.03 |
| 19-131 | 16 | 0.015 |
| 19-51 | 24 | 0.015 |
| 19-132 | 24 | 0.03 |
| 19-133 | 24 | 0.03 |
| 19-134 | 32 | 0.03 |
| GEOMEAN | 7.046 | 0.026 |

**Supplementary Table 5.** MGX MICs (mg/L) for *C. auris* isolates with high azole MICs

| *C. auris* | FLC | VRC | ITC | ISA | POS | MGX |
| --- | --- | --- | --- | --- | --- | --- |
| 18-11 | 256 | 4 | 0.5 | 1 | 0.25 | 0.03 |
| 18-59 | 256 | 4 | 0.5 | 1 | 0.5 | 0.015 |
| 19-23 | 256 | 4 | 0.5 | 0.5 | 0.12 | 0.008 |
| 19-50 | 256 | 4 | 1 | 1 | 0.5 | 0.015 |
| 19-60 | 256 | 4 | 1 | 1 | 1 | 0.015 |
| 19-84 | 256 | 4 | 1 | 1 | 0.25 | 0.03 |
| 19-93 | 256 | 4 | 1 | 1 | 0.5 | 0.03 |
| 19-94 | 256 | 4 | 0.5 | 0.25 | 0.25 | 0.008 |
| 19-95 | 256 | 4 | 1 | 1 | 0.5 | 0.03 |
| 19-96 | 256 | 4 | 1 | 1 | 0.5 | 0.03 |
| 18-57 | 256 | 2 | 1 | 2 | 0.25 | 0.015 |
| 18-58 | 256 | 2 | 1 | 2 | 0.5 | 0.03 |
| GEOMEAN | 256.000 | 3.564 | 0.794 | 0.944 | 0.373 | 0.020 |

**Supplementary Table 6.**
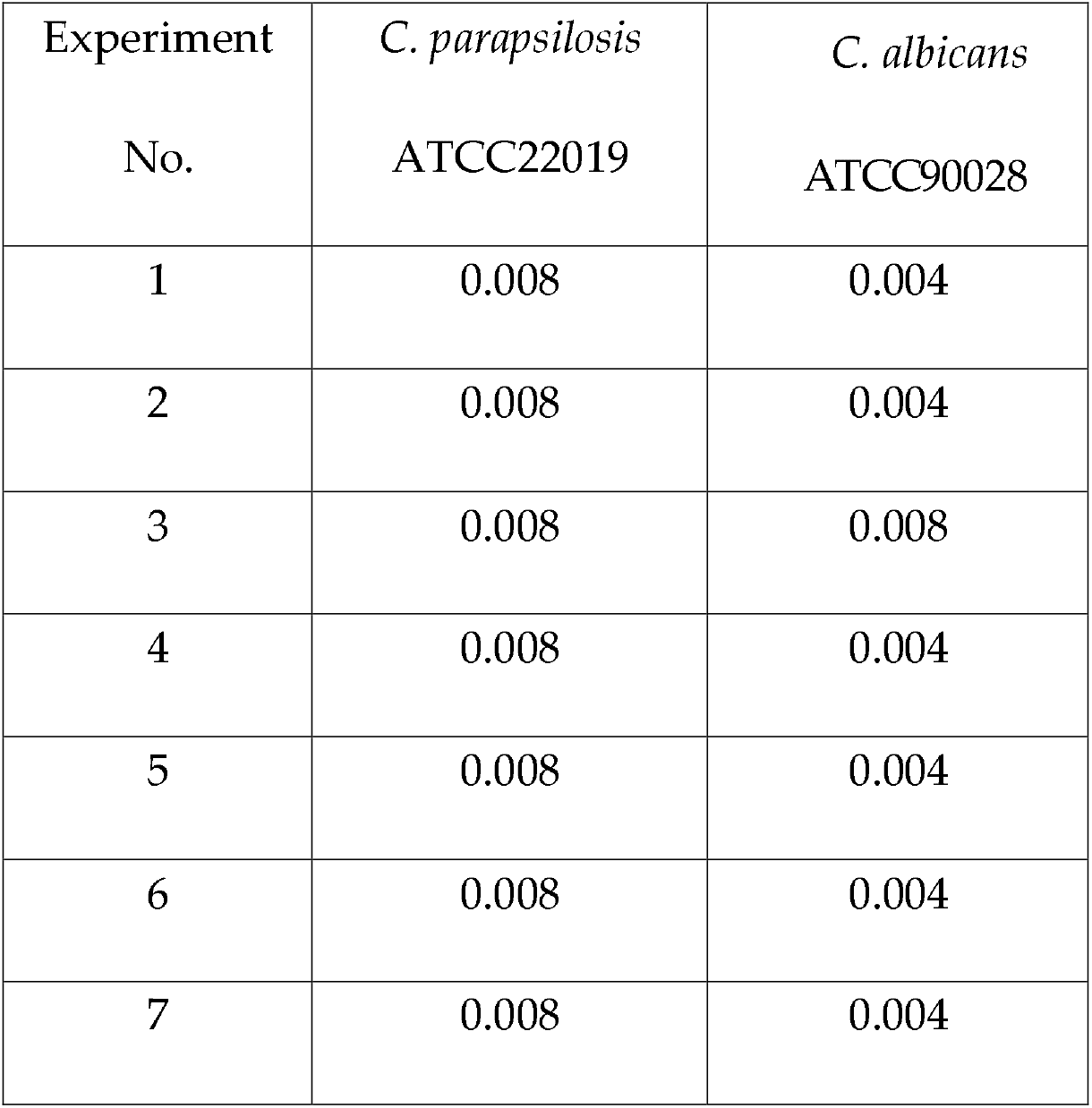
MGX MIC (mg/L) values for two *Candida* quality control (QC) strains.

